# Vulnerability of Elevation-Restricted Endemic Birds of the Cordillera de Talamanca (Costa Rica and Panama) to Climate Change

**DOI:** 10.1101/2020.09.11.293134

**Authors:** Zhen Liu, Luis Sandoval, Lauren Sherman, Andrew Wilson

## Abstract

Animals endemic to tropical mountains are known to be especially vulnerable to climate change. The Cordillera de Talamanca (Costa Rica and Panama) is a geographically isolated mountain chain and global biodiversity hotspot, home to more than 50 endemic bird species. We used eBird community science observations to predict the distributions of a suite of 48 of these endemic birds in 2006-2015, and in 2070, under four climate change scenarios. Species distributions were predicted using program Maxent, incorporating elevation, satellite derived habitat data, and WorldClim climate variables. Model fit, as assessed by Area under the Receiver Operator Curve (AUC) was very high for most species, ranging from 0.877 to 0.992 (mean of 0.94). We found that most species are predicted to undergo range contractions by 2070, with a mean of 15% under modest climate change (RCP 2.6) up to a mean of 40% under more severe climate change (RCP 8.5). Most of the current ranges of these species are within existing protected areas (average of 59% in 2006-2015), and with prospective range contractions, the importance of these protected areas is forecast to increase. We suggest that these predicted range declines should elevate conservation concerns for this suite of species, and vigilance, in the form of better population monitoring, is urgently needed.

## INTRODUCTION

The global average air temperature has risen by more than 1°C since pre-industrial times (Schurer et al. 2017), changing at a rate faster than many species can adapt (Abolafya et al. 2013). By the year 2100, the surface of the earth could have warmed by 3.5°C, which could result in 600 to 900 extinctions of land bird species, with 89% of those in the tropics (Şekercioğlu et al. 2012). Species found at elevations of more than 500 m are more sensitive to warming temperatures because they have low basal metabolic rates and experience limited temperature variation (Şekercioğlu et al. 2008). Upward shifts could result in an “escalator to extinction” for some high elevation species, which simply run out of room as the area of suitable habitat diminishes (Freeman et al. 2018).

On average, birds have been documented to be shifting their ranges upslope by 11 meters per decade (Chen et al. 2011), but shifts are larger for birds found in the tropics than those from temperate regions (Freeman and Class Freeman 2014). The causal mechanism for the stronger response of tropical montane bird species to climate change is not well understood, but it could be due to changes in prey abundance and adaptation to narrow temperature regimes (Freeman et al. 2018). There are various drivers of these potential range shifts, including internal species traits (e.g. time delay in species’ response, individualistic physiological constraints), and external drivers of change (e.g. multispecies interactions, non-climatic factors) (Chen et al. 2011). Furthermore, the effects of climate change on tropical birds will be exacerbated by the synergies of climate change and other threats such as diseases, invasive species, hunting, and habitat loss (Şekercioğlu et al. 2012).

Despite these predictions, there are some uncertainties. Birds have been shown to be capable of closely tracking warming temperatures (Devictor et al. 2008) and because they are mobile, it is relatively easy for them to move upslope compared with less mobile organisms. Why then would they still face the possibility of going extinct? While birds could potentially shift their range, their food resources and habitat might not be able to shift within a short period of time. For example, synchronizing their reproductive cycle with the emergence of food has been identified as a serious challenge for insectivorous birds in the face of climate change (Charmantier et al. 2008). The effects of climate change on some ecosystems are very specific. One example is the cloud forests of Central America. Cloud forests are montane forests that have permanently moist climates due to frequent low-cloud, and are among the most biodiverse habitats globally (Karmalkar et al. 2008). According to a global survey of elevation-restricted birds, over 20% of species are found primarily in cloud forest, and many of these species are endangered (Foster 2001).

Southern Central America is one of the most biodiverse places in the world (Mittermeier et al. 1998), with more breeding bird species than the United States and Canada combined (Stiles and Skutch 1989, Ridgely and Gwynne 1992). Many bird species found in Central America are endemic to the region (Stiles and Skutch 1989, Ridgely and Gwynne 1992). Of those range-restricted species, many are found at high elevations in the Cordillera de Talamanca, or on isolated volcanoes in the north of Costa Rica, within the geographical area called the Talamanca Montane Forests (Stiles and Skutch 1989, Ridgely and Gwynne 1992). These species, with small geographic ranges and sometimes narrow elevational ranges, are vulnerable to future decline, as climate change reduces their prospective habitats. Latitudinal shifts away from the equator are not an adaptation option for these species, because forested areas to the immediate north of this region are at low elevation for several hundred kilometers (Chavarría-Pizarro et al. 2010, Barrantes et al. 2011).

In this study, we the future geographic ranges of 48 elevation-restricted birds endemic to Cordillera de Talamanca and surrounding areas, when faced with predicted climate change. While our study region focusses on the Cordillera de Talamanca, which reaches a maximum elevation of 3,819 meters at Cerro Chirripó, we include the whole of Costa Rica and Panama west of 70° west (Figure 1), to encompass the entire geographic ranges of the focal endemic birds (Barrantes 2009). Almost all these species are currently listed as of “Least Concern” on the Red List of International Union for Conservation of Nature, with the only exception being the Red-fronted Parrotlet *Touit costaricensis*, which is listed as “Vulnerable” (Table 1). All species are endemic to Costa Rica and Panama, and have global range sizes that average less than 10,000 km^2^ (BirdLife International. 2019). Most of the study species are forest obligates, and hence largely restricted to the Talamancan montane forests ecosystem, but some of the most elevationally restricted are found in the páramo shrub and high elevation grasslands that are found above 3,000 meters (Stiles and Skutch 1989, Ridgely and Gwynne 1992, Barrantes 2009, Barrantes et al. 2011).

**Figure 1.**
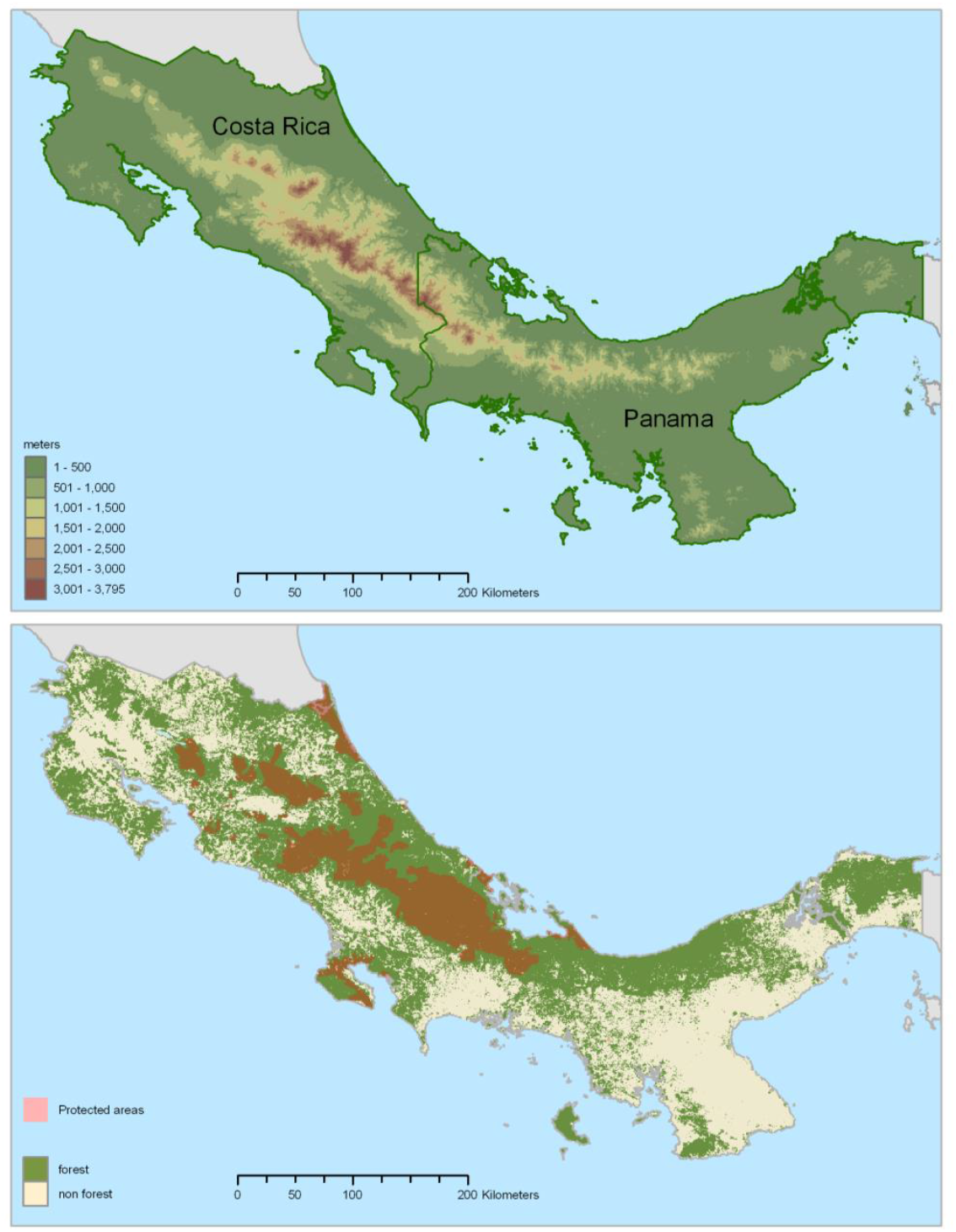
Elevation within the study area of Costa Rica and western Panama (top), and forested (MODIS land cover) and protected areas (bottom) from the World Database on Protected Areas (IUCN 2017).

**Table 1.** List of species included in the analysis, with IUCN Red List categorization. * Species that conduct altitudinal migration

| English name | Scientific name | IUCN Red List category | eBird locations used in analysis |
| --- | --- | --- | --- |
| Black Guan* | *Chamaepetes unicolor* | Least Concern | 271 |
| Black-breasted Wood-Quail | *Odontophorus leucolaemus* | Least Concern | 131 |
| Buff-fronted Quail-Dove | *Zentrygon costaricensis* | Least Concern | 80 |
| Chiriqui Quail-Dove | *Zentrygon chiriquensis* | Least Concern | 109 |
| Dusky Nightjar | *Antrostomus saturatus* | Least Concern | 94 |
| Talamanca Hummingbird | *Eugenes spectabilis* | † | 197 |
| Fiery-throated Hummingbird | *Panterpe insignis* | Least Concern | 227 |
| White-bellied Mountain-gem | *Lampornis hemileucus* | Least Concern | 114 |
| Magenta-throated Woodstar* | *Calliphlox bryantae* | Least Concern | 140 |
| Volcano Hummingbird | *Selasphorus flammula* | Least Concern | 205 |
| Scintillant Hummingbird* | *Selasphorus scintilla* | Least Concern | 295 |
| Black-bellied Hummingbird | *Eupherusa nigriventris* | Least Concern | 125 |
| White-tailed Emerald | *Elvira chionura* | Least Concern | 141 |
| Coppery-headed Emerald | *Elvira cupreiceps* | Least Concern | 164 |
| Costa Rican Pygmy Owl | *Glaucidium costaricanum* | Least Concern | 67 |
| Prong-billed Barbet | *Semnornis frantzii* | Least Concern | 217 |
| Sulphur-winged Parakeet | *Pyrrhura hoffmanni* | Least Concern | 259 |
| Red-fronted Parrotlet* | *Touit costaricensis* | Vulnerable | 83 |
| Silvery-fronted Tapaculo | *Scytalopus argentifrons* | Least Concern | 217 |
| Streak-breasted Treehunter | *Thripadectes rufobrunneus* | Least Concern | 144 |
| Ruddy Treerunner | *Margarornis rubiginosus* | Least Concern | 218 |
| Golden-bellied Flycatcher | *Myiodynastes hemichrysus* | Least Concern | 231 |
| Dark Pewee | *Contopus lugubris* | Least Concern | 235 |
| Ochraceous Pewee | *Contopus ochraceus* | Least Concern | 35 |
| Black-capped Flycatcher | *Empidonax atriceps* | Least Concern | 126 |
| Yellow-winged Vireo | *Vireo carmioli* | Least Concern | 219 |
| Silvery-throated Jay | *Cyanolyca argentigula* | Least Concern | 38 |
| Ochraceous Wren | *Troglodytes ochraceus* | Least Concern | 330 |
| Timberline Wren | *Thryorchilus browni* | Least Concern | 49 |
| Black-faced Solitaire | *Myadestes melanops* | Least Concern | 436 |
| Black-billed Nightingale-Thrush | *Catharus gracilirostris* | Least Concern | 162 |
| Sooty Thrush | *Turdus nigrescens* | Least Concern | 153 |
| Black-and-yellow Silky-flycatcher | *Phainoptila melanoxantha* | Least Concern | 208 |
| Long-tailed Silky-flycatcher | *Ptiliogonys caudatus* | Least Concern | 302 |
| Golden-browed Chlorophonia* | *Chlorophonia callophrys* | Least Concern | 358 |
| Sooty-capped Chlorospingus | *Chlorospingus pileatus* | Least Concern | 301 |
| Sooty-faced Finch | *Arremon crassirostris* | Least Concern | 87 |
| Volcano Junco | *Junco vulcani* | Least Concern | 43 |
| Large-footed Finch | *Pezopetes capitalis* | Least Concern | 149 |
| Yellow-thighed Brushfinch | *Atlapetes tibialis* | Least Concern | 283 |
| Wrenthrush | *Zeledonia coronata* | Least Concern | 116 |
| Flame-throated Warbler | *Oreothlypis gutturalis* | Least Concern | 245 |
| Black-cheeked Warbler | *Basileuterus melanogenys* | Least Concern | 242 |
| Collared Redstart | *Myioborus torquatus* | Least Concern | 322 |
| Black-thighed Grosbeak | *Pheucticus tibialis* | Least Concern | 206 |
| Spangle-cheeked Tanager* | *Tangara dowii* | Least Concern | 294 |
| Peg-billed Finch | *Acanthidops bairdi* | Least Concern | 78 |
| Slaty Flowerpiercer | *Diglossa plumbea* | Least Concern | 336 |

†not assessed by IUCN as of July 2020 (Partida-Lara and Enríquez 2020)

## METHODS

We used program Maxent (Phillips et al. 2006) to predict baseline (2006-2015), and future (2070) species distributions under the four climate change scenarios of the IPCC’s AR5 Greenhouse Gas Concentration Pathways (IPCC 2014).

### Data Sources

To predict baseline species distributions, we used data from the Cornell Lab of Ornithology’s eBird program—a community science database of bird observations compiled from checklists submitted through an online portal (Sullivan et al. 2009). eBird data are verified by both model-based automated filters and a network of voluntary reviewers (Kelling et al. 2011). We used bird observations for the 10-year period 2006-2015 to estimate baseline distributions. To reduce the effects of sampling bias we thinned locations, eliminating duplicate locations less than 2 km apart, using R package SPthin (Aiello-Lammens et al. 2015). We choose 2 km for thinning because many of the species have small geographic distributions and thinning by larger distances would result in a greatly diminished sample for some species. Following thinning there were between 35 and 436 locations for the 48 study species (Table 1), with a mean of 193 locations per species. The Talamanca Hummingbird *Eugenes spectabilis* was added to the analysis following it’s split from the Rivoli’s Hummingbird *Eugenes fulgens* in 2017 (Chesser et al. 2017)

Environment data layers include elevation (SRTM DEM; (Jarvis et al. 2008)), land cover, ecosystem typing, and climate. We used the Model Builder in ArcMap 10 (ESRI 2011) to clip each environmental raster to the study area to standardized projection, extent and cell size (90 x 90 m) and convert them to ASCII format for use in Maxent.

Land cover type was derived from Moderate Resolution Imaging Spectroradiometer (MODIS) satellite imagery obtained in 2001-2012 (Friedl et al. 2010). These images are based on International Geosphere–Biosphere Programme (IGBP) ecosystem classes (Strahler and Muller 1999), which do not differentiate between types of broadleaf evergreen forest, which is the dominant forest type in our study area, comprising 50.5% of the land surface. To provide a more nuanced representation of forest types we also included an ecosystem typing based on the Central American Ecosystems Map from year 2000 (Meyrat et al. 2002) reclassified into 18 ecosystem types (Appendix 1).

We obtained raster layers of monthly climate data from Version 2 of the WorldClim database (Fick and Hijmans 2017). Future climate projections were from the *HadGEM2-ES* global climate model. Based on previous studies (Buermann et al. 2008, Velásquez-Tibatá et al. 2013, Avalos and Hernández 2015, Ortega-Huerta and Vega-Rivera 2017) we selected seven climate variables for inclusion in the models, representing both temperature and precipitation metrics:

BIO1 = Annual Mean Temperature
BIO4 = Temperature Seasonality (standard deviation *100)
BIO5 = Max Temperature of Warmest Month
BIO6 = Min Temperature of Coldest Month
BIO7 = Temperature Annual Range (BIO5-BIO6)
BIO12 = Annual Precipitation
BIO15 = Precipitation Seasonality (Coefficient of Variation)

Climate data were for a baseline period of 1970 to 2000, and the four future scenarios for the year 2070: RCP 2.6, RCP 4.5, RCP 6.0, and RCP 8.5. RCP 2.6 assumes that global greenhouse gas emissions peak between 2010 and 2020, and then decline; RCP 4.5 and 6.0 assume that emissions continue to rise, with peak emissions occurring in 2040 and 2080 respectively, while RCP 8.5 assumes emissions continue to rise throughout the 21^st^ Century. Hence climate change would be modest under RCP 2.6, more severe under 4.5 and 6.0, and most severe under RCP 8.5 (Meinshausen et al. 2011). As suggested by other authors (McCollum et al. 2013, Lyra et al. 2017), we assume that the RCP 8.5 is the most likely scenario, and that RCP 6.0 would be a more moderate scenario that could occur if the global community takes major steps to reduce emissions in coming decades. All temperatures were in x10 degrees Celsius. Climate data were aligned and downscaled to 90 m x 90 m cells to ensure compatibility with landscape data.

### Species Distribution Models

We ran Maxent with a regularization multiplier of 2, to avoid overfitting (Radosavljevic and Anderson 2014). We tested model performance using Area under the Receiver Operator Curve (AUC) based on a reserved random selection of 20% of sample locations as the test points, and ran a total of 10 replicates, with up to 1500 iterations for each species. We used the Maxent clamping procedure to constrain the range of projected values to the range of the background values used to train the model (Phillips et al. 2006).

To produce maps of estimated area of occupancy in 2006-2015 and for each climate change scenario in 2070, we classified the model output (probability of suitability in each cell, ranging from 0 to 1) into two classes, suitable (0) and unsuitability (1). There are many methods available for classifying probability data into two discrete classes (Escalante et al. 2013). We used the “Equal test sensitivity and specificity” (ETSS) complementary log-log threshold, which balances errors of omission and commission (Phillips et al. 2006). The threshold was applied to the Maxent output maps in ArcGIS 10, and the areas with probabilities above the ETSS threshold were assumed suitable, i.e. to be within the species’ baseline range.

### Protected Areas Outlook

Based on the predicted area of occupancy we calculated how much of the 2006-2015 and 2070 ranges were within protected areas. We overlaid a map of currently protected areas, taken from the World Database on Protected Areas (WDPA, November 2019 update; UNEP-WCMC and IUCN 2019), with raster data of the projected species ranges in Program R. From these data we were able to calculate the percentage of range for each species that remained in protected areas under each projected climate outcome.

## RESULTS

We found that model fit was excellent for all 48 endemic species analyzed, with a mean AUC of 0.941 (Table 2). Only one species had an AUC of less than 0.9, the Red-fronted Parrotlet (0.877; Table 2). AUC values of greater than 0.9 indicate excellent model accuracy (Swets 1988). These high AUC value indicates that the current distributions of these species can be accurately predicted based on the seven environmental variables selected for our study. Estimated baseline ranges and range sizes were similar to species distribution maps curated by BirdLife international for all of our study species (e.g. for Volcano Junco *Junco vulcani*; Figure 2). However, there was a general pattern of over-estimation of range size for species with the most restricted ranges, but under-estimation for the more widespread species (Figure 3). Our baseline range sizes were larger than birdlife ranges by a mean of 8%, varying from 59% smaller for the Black-faced Solitaire *Myadestes melanops*, up to 175% larger for the Sooty Thrush *Turdus nigrescens* (Table 2).

**Table 2.** Area under the Curve (AUC) scores for our Maxent models for all 48 bird species, baseline range size estimates from Birdlife International maps and Maxent models based on eBird data from 2006-15, and predicted future range sizes and range changes under 4 climate change scenarios.

| Scientific name | AUC | Estimated range size in 1,000s km<sup>2</sup> (percent change from 2006-2015) |  |  |  |  |  |
| --- | --- | --- | --- | --- | --- | --- | --- |
|  |  | Our study |  |  |  |  |  |
|  |  | Baseline | 2070 under 4 RCPs |  |  |  |  |
|  |  | BirdLife | 2006-15 | 2.6 | 4.5 | 6 | 8.5 |
| *Chamaepetes unicolor* | 0.925 | 15.57 | 12.8 | 15.5 (+21.1) | 14.1 (+10.2) | 13.7 (+7.0) | 13.7 (+7.0) |
| *Odontophorus leucolaemus* | 0.935 | 9.48 | 11.4 | 11.5 (+0.9) | 7.7 (-32.5) | 6.9 (-39.5) | 6.1 (-46.5) |
| *Zentrygon costaricensis* | 0.934 | 16.52 | 11.9 | 9.4 (-21.0) | 7.2 (-39.5) | 6.1 (-48.7) | 4.6 (-61.3) |
| *Zentrygon chiriquensis* | 0.929 | 16.38 | 13.6 | 11.1 (-18.4) | 8.5 (-37.5) | 6.5 (-52.2) | 5.8 (-57.4) |
| *Antrostomus saturatus* | 0.961 | 5.38 | 8.5 | 6.2 (-27.1) | 5.3 (-37.6) | 5.2 (-38.8) | 4 (-52.9) |
| *Eugenes spectabilis* | 0.956 | * | 9.1 | 8.3 (-8.8) | 8.1 (-11.0) | 7.8 (-14.3) | 5.7 (-37.4) |
| *Panterpe insignis* | 0.957 | 8.27 | 8.7 | 6.8 (-21.8) | 5.1 (-41.4) | 4.1 (-52.9) | 3.4 (-60.9) |
| *Lampornis hemileucus* | 0.929 | 9.35 | 12.5 | 15 (+20) | 12.9 (+3.2) | 13.9 (+11.2) | 13.2 (+5.6) |
| *Calliphlox bryantae* | 0.915 | 9.19 | 13.6 | 11 (-19.1) | 9.3 (-31.6) | 8.5 (-37.5) | 6.9 (-49.3) |
| *Selasphorus flammula* | 0.96 | 4.6 | 8.2 | 6.1 (-25.6) | 5.7 (-30.5) | 5.5 (-32.9) | 4.5 (-45.1) |
| *Selasphorus scintilla* | 0.941 | 9.42 | 9.8 | 6.8 (-30.6) | 6.5 (-33.7) | 5.9 (-39.8) | 4.8 (-51.0) |
| *Eupherusa nigriventris* | 0.91 | 14.34 | 13.7 | 9.8 (-28.5) | 6.7 (-51.1) | 6.8 (-50.4) | 3.9 (-71.5) |
| *Elvira chionura* | 0.941 | 9.6 | 10 | 6.7 (-33.0) | 8.5 (-15.0) | 8.6 (-14.0) | 8.8 (-12.0) |
| *Elvira cupreiceps* | 0.941 | 5.22 | 9.8 | 14.8 (+51.0) | 9.2 (-6.1) | 9.3 (-5.1) | 8.8 (-10.2) |
| *Glaucidium costaricanum* | 0.959 | 10.6 | 10.4 | 5.8 (-44.5) | 4.8 (-53.8) | 4.1 (-61.2) | 3.7 (-64.3) |
| *Semnornis frantzii* | 0.928 | 15.87 | 11.1 | 11.2 (0.9) | 8.7 (-21.6) | 8.8 (-20.7) | 8.2 (-26.1) |
| *Pyrrhura hoffmanni* | 0.921 | 8 | 12.7 | 6.4 (-49.6) | 5.9 (-53.5) | 5.2 (-59.1) | 2.5 (-80.3) |
| *Touit costaricensis* | 0.877 | 16.63 | 19.3 | 13.4 (-30.6) | 5.5 (-71.5) | 5.7 (-70.5) | 2.6 (-86.5) |
| *Scytalopus argentifrons* | 0.935 | 15.55 | 11.6 | 10.3 (-11.2) | 8.5 (-26.7) | 8.2 (-29.3) | 6 (-48.3) |
| *Thripadectes rufobrunneus* | 0.931 | 17.59 | 12.1 | 12 (-0.8) | 9.7 (-19.8) | 9.6 (-20.7) | 8.9 (-26.4) |
| *Margarornis rubiginosus* | 0.948 | 5.6 | 9.6 | 5.8 (-39.6) | 4.2 (-56.3) | 3.7 (-61.5) | 2.8 (-70.8) |
| *Myiodynastes hemichrysus* | 0.903 | 16.61 | 14.7 | 11.7 (-20.4) | 6.2 (-57.8) | 5.7 (-61.2) | 3 (-79.6) |
| *Contopus lugubris* | 0.92 | 8.2 | 12.2 | 7.2 (-41.0) | 5.9 (-51.6) | 5.1 (-58.2) | 2.8 (-77.0) |

| Scientific name | AUC | Estimated range size in 1,000s km<sup>2</sup> (percent change from 2006-2015) |  |  |  |  |  |  |  |  |  |
| --- | --- | --- | --- | --- | --- | --- | --- | --- | --- | --- | --- |
| *Contopus ochraceus* | 0.976 | 3.14 | 5.1 | 3 | (-41.2) | 2.6 | (-49.0) | 2.4 | (-52.9) | 1.7 | (-66.7) |
| *Empidonax atriceps* | 0.978 | 3.66 | 5.1 | 4.7 | (-7.8) | 4.3 | (-15.7) | 4.3 | (-15.7) | 4.1 | (-19.6) |
| *Vireo carmioli* | 0.95 | 4.19 | 9.5 | 6.9 | (-27.4) | 6.6 | (-30.5) | 6.3 | (-33.7) | 5.1 | (-46.3) |
| *Cyanolyca argentigula* | 0.986 | 3.92 | 4.3 | 2.3 | (-46.5) | 2.1 | (-51.2) | 1.9 | (-55.8) | 1.2 | (-72.1) |
| *Troglodytes ochraceus* | 0.914 | 17.33 | 13.3 | 17.1 | (+28.6) | 14.4 | (+8.3) | 14.5 | (+9.0) | 14 | (+5.3) |
| *Thryorchilus browni* | 0.99 | 1.49 | 2.7 | 2.1 | (-22.2) | 2.1 | (-22.2) | 2.1 | (-22.2) | 1.5 | (-44.4) |
| *Myadestes melanops* | 0.913 | 31.92 | 13 | 10.7 | (-17.7) | 7.6 | (-41.5) | 7.1 | (-45.4) | 4.7 | (-63.8) |
| *Catharus gracilirostris* | 0.972 | 3.83 | 6 | 5.3 | (-11.7) | 5.1 | (-15.0) | 5 | (-16.7) | 4.7 | (-21.7) |
| *Turdus nigrescens* | 0.973 | 2.07 | 5.7 | 7.6 | (+33.3) | 7.5 | (+31.6) | 8.6 | (+50.9) | 8.6 | (+50.9) |
| *Phainoptila melanoxantha* | 0.945 | 6.57 | 9.7 | 8.5 | (-12.4) | 7.4 | (-23.7) | 7.4 | (-23.7) | 6 | (-38.1) |
| *Ptiliogonys caudatus* | 0.941 | 9.54 | 9.6 | 7.4 | (-22.9) | 7 | (-27.1) | 6.7 | (-30.2) | 5.5 | (-42.7) |
| *Chlorophonia callophrys* | 0.917 | 17.43 | 13 | 11.9 | (-8.5) | 6.5 | (-50) | 9.2 | (-29.2) | 7.4 | (-43.1) |
| *Chlorospingus pileatus* | 0.948 | 3.94 | 9.8 | 8.3 | (-15.3) | 7.4 | (-24.5) | 7.2 | (-26.5) | 6.3 | (-35.7) |
| *Arremon crassirostris* | 0.931 | 9.5 | 12 | 13 | (+8.3) | 6.3 | (-47.5) | 8.1 | (-32.5) | 8.6 | (-28.3) |
| *Junco vulcani* | 0.992 | 1.04 | 2.1 | 1.7 | (-19.0) | 1.6 | (-23.8) | 1.5 | (-28.6) | 1.5 | (-28.6) |
| *Pezopetes capitalis* | 0.972 | 3.4 | 6.2 | 5.2 | (-16.1) | 5.1 | (-17.7) | 4.9 | (-21.0) | 4.8 | (-22.6) |
| *Atlapetes tibialis* | 0.94 | 6.56 | 10 | 9.2 | (-8.0) | 8.1 | (-19.0) | 7.7 | (-23.0) | 7.3 | (-27.0) |
| *Zeledonia coronata* | 0.956 | 5.17 | 9 | 7.3 | (-18.9) | 6.3 | (-30.0) | 5.6 | (-37.8) | 5.9 | (-34.4) |
| *Oreothlypis gutturalis* | 0.952 | 3.91 | 8.7 | 6.7 | (-23.0) | 6.7 | (-23.0) | 6.5 | (-25.3) | 5.5 | (-36.8) |
| *Basileuterus melanogenys* | 0.946 | 4.83 | 9.8 | 8 | (-18.4) | 7.8 | (-20.4) | 7.8 | (-20.4) | 6.6 | (-32.7) |
| *Myioborus torquatus* | 0.94 | 5.26 | 10.5 | 8.8 | (-16.2) | 7.4 | (-29.5) | 7.2 | (-31.4) | 5.8 | (-44.8) |
| *Pheucticus tibialis* | 0.908 | 19.6 | 15.2 | 17.1 | (+12.5) | 13.6 | (-10.5) | 15.2 | (0) | 13.7 | (-9.9) |
| *Tangara dowii* | 0.923 | 12.08 | 12.1 | 10.5 | (-13.2) | 7.3 | (-39.7) | 7.1 | (-41.3) | 4.7 | (-61.2) |
| *Acanthidops bairdi* | 0.967 | 4.74 | 8.4 | 7 | (-16.7) | 6.6 | (-21.4) | 6.3 | (-25.0) | 5.7 | (-32.1) |
| *Diglossa plumbea* | 0.938 | 8.93 | 11.1 | 10 | (-9.9) | 9 | (-18.9) | 9 | (-18.9) | 8 | (-27.9) |

**Figure 2.**
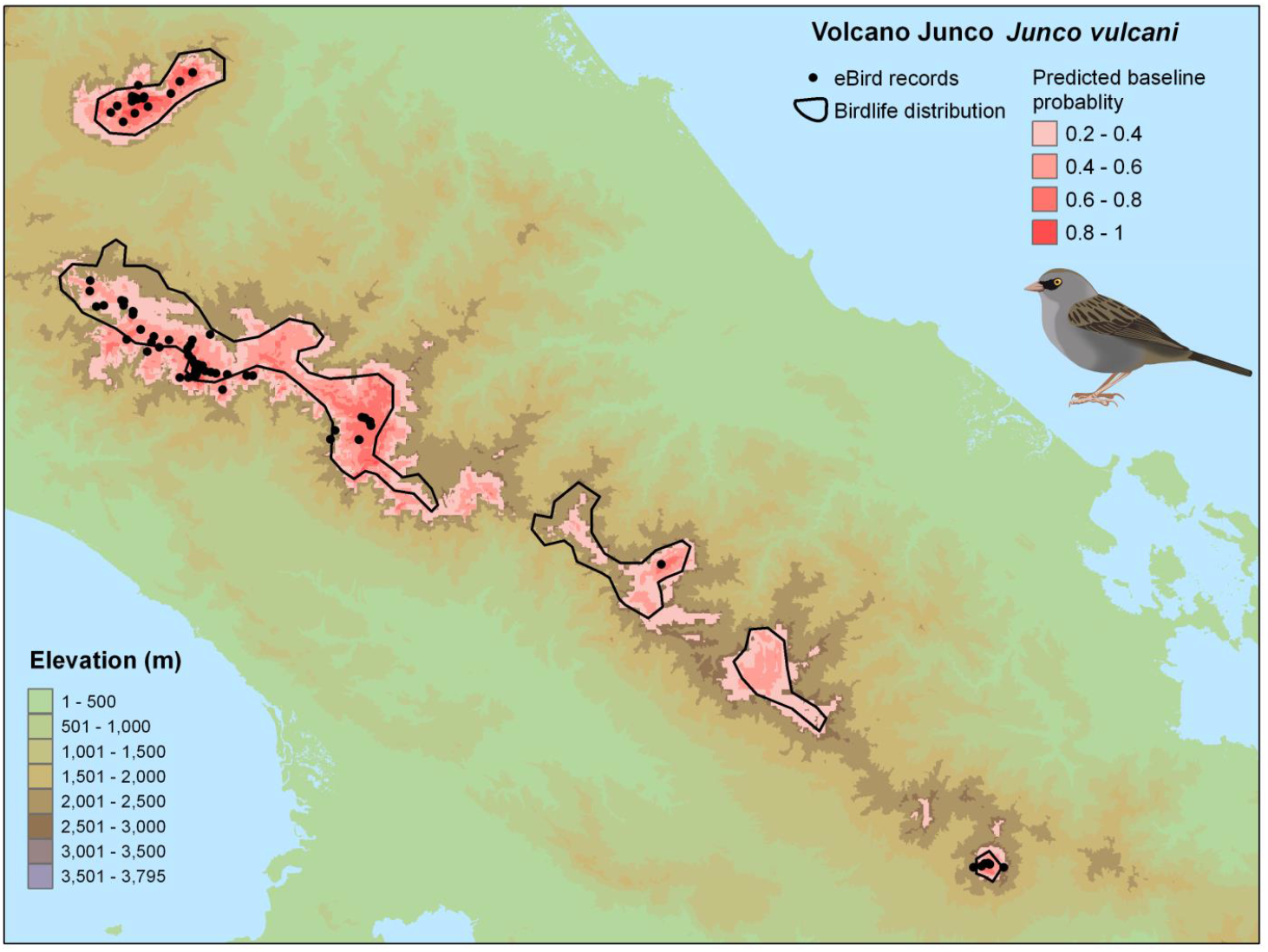
Current distribution (BirdLife International. 2019) (BirdLife International. 2019), locations of eBird records (post-thinning) and our baseline (2006-2015) predicted range of the Volcano Junco *Junco vulcani*

**Figure 3.**
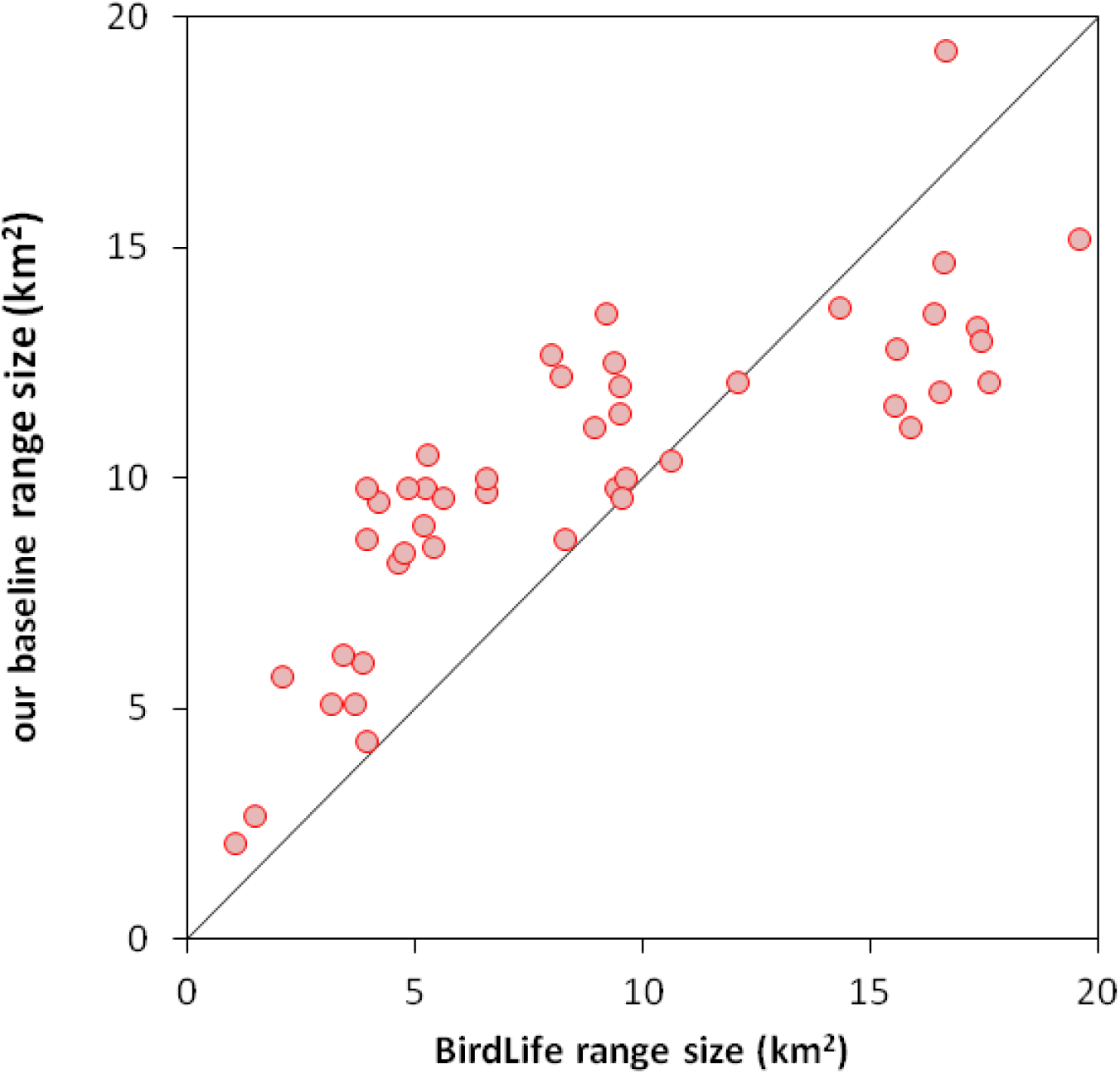
Comparison between estimated range sizes from Birdlife (BirdLife International 2019) and our baseline (2006-2015) estimate.

Our models predict that under RCP 2.6, 39 of the 48 species would undergo range contractions, with a mean range decline of 14.4% (Figure 4), but no species were predicted to undergo range declines of more than 50%. Under the intermediate scenarios, the number of species predicted to undergo range declines was 44 and 43 under RCP 4.5 and RCP 6.0 respectively. However, the number of species predicted to undergo range reductions of more than 50% was 9 and 11 under RCP 4.5 and RCP 6.0 respectively. Under the more extreme RCP 8.5 scenarios, 45 species were predicted to undergo range declines, 16 of them showing predicted range declines of more than 50%.

**Figure 4.**
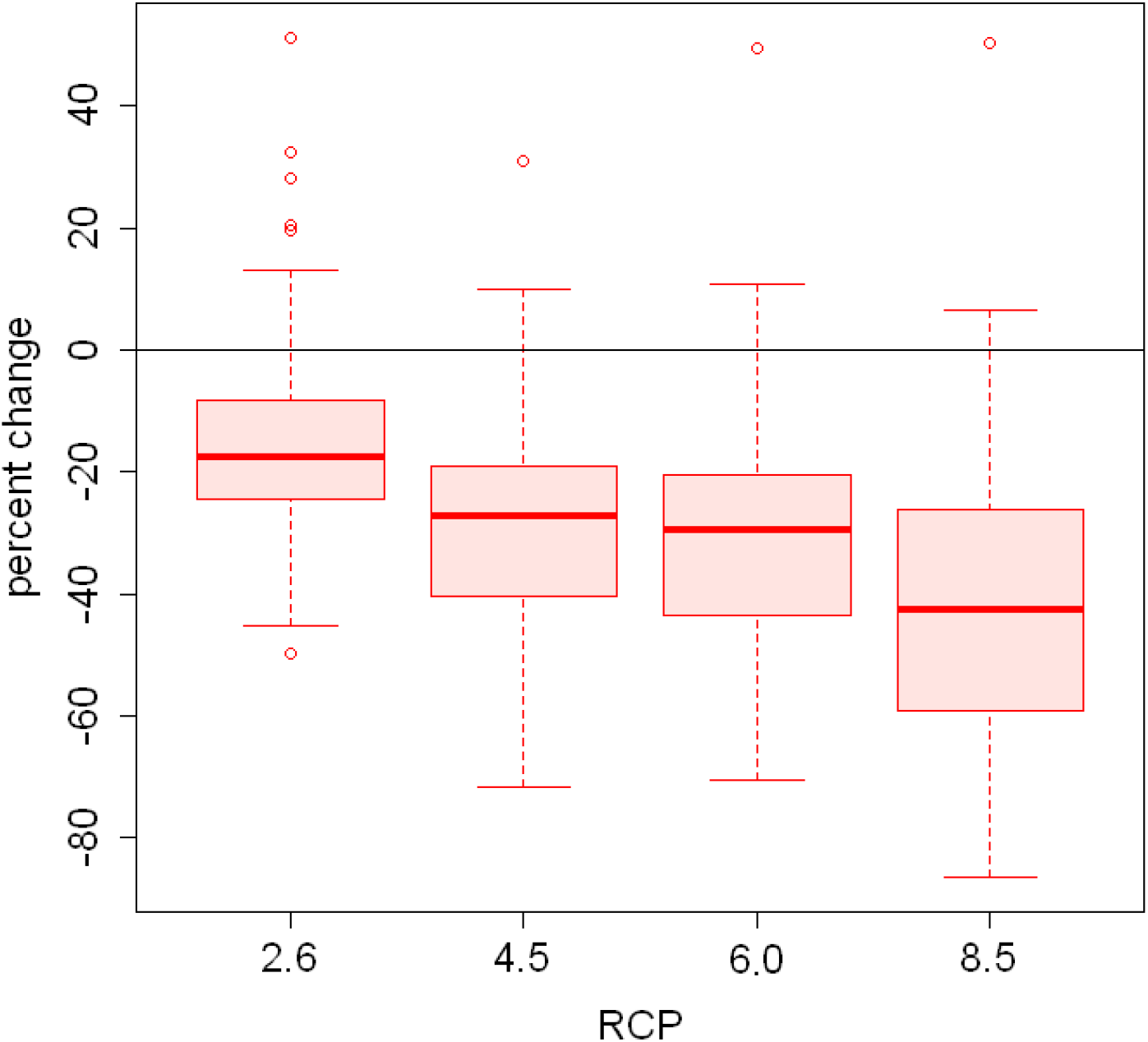
Predicted percent changes in range size by 2070 for the 48 species under the four IPCC AR5 Greenhouse Gas Concentration Pathways (IPCC 2014). Circles indicate outliers.

Upslope range contractions could result in a loss of these endemic birds from some mid-elevation protected areas. The Collared Redstart *Myioborus torquatus*, for example—a species whose predicted range contractions were close to the average among the 48 species—is predicted to be lost from protected areas, such as the cloud forests of Monteverde and Arenal Volcano National Park (Figure 5). We estimate that on average 58.9% of species’ ranges were in protected areas in 2006–2015 (Figure 6). Overall, the average percentages of ranges in protected areas is predicted to increase slightly—to 60.9% under RCP 2.6, 64.0% under RCP 4.5, 63.9% under RCP 6.0 and 67.2% under RCP 8.5. For most species, more than 50% of their range is predicted to be protected now and in the future, exceptions being White-tailed Emerald *Elvira chionura* (39.6% baseline, 37.4–40.4% future), Coppery-headed Emerald *Elvira cupreiceps* (41.8% baseline, 40.0–46.7% future), and Red-fronted Parrotlet (44.5% baseline, 45.4–53.1% future). At the other extreme, species with the highest percentage of ranges protected include páramo or montane forest specialists, such as Volcano Junco (83.7% baseline, 84.8-85.0% future), Timberline Wren (80.4% now, 84.4–85.0% future), and Silvery-throated Jay (78.5% baseline, 86.7–89.3% future).

**Figure 5.**
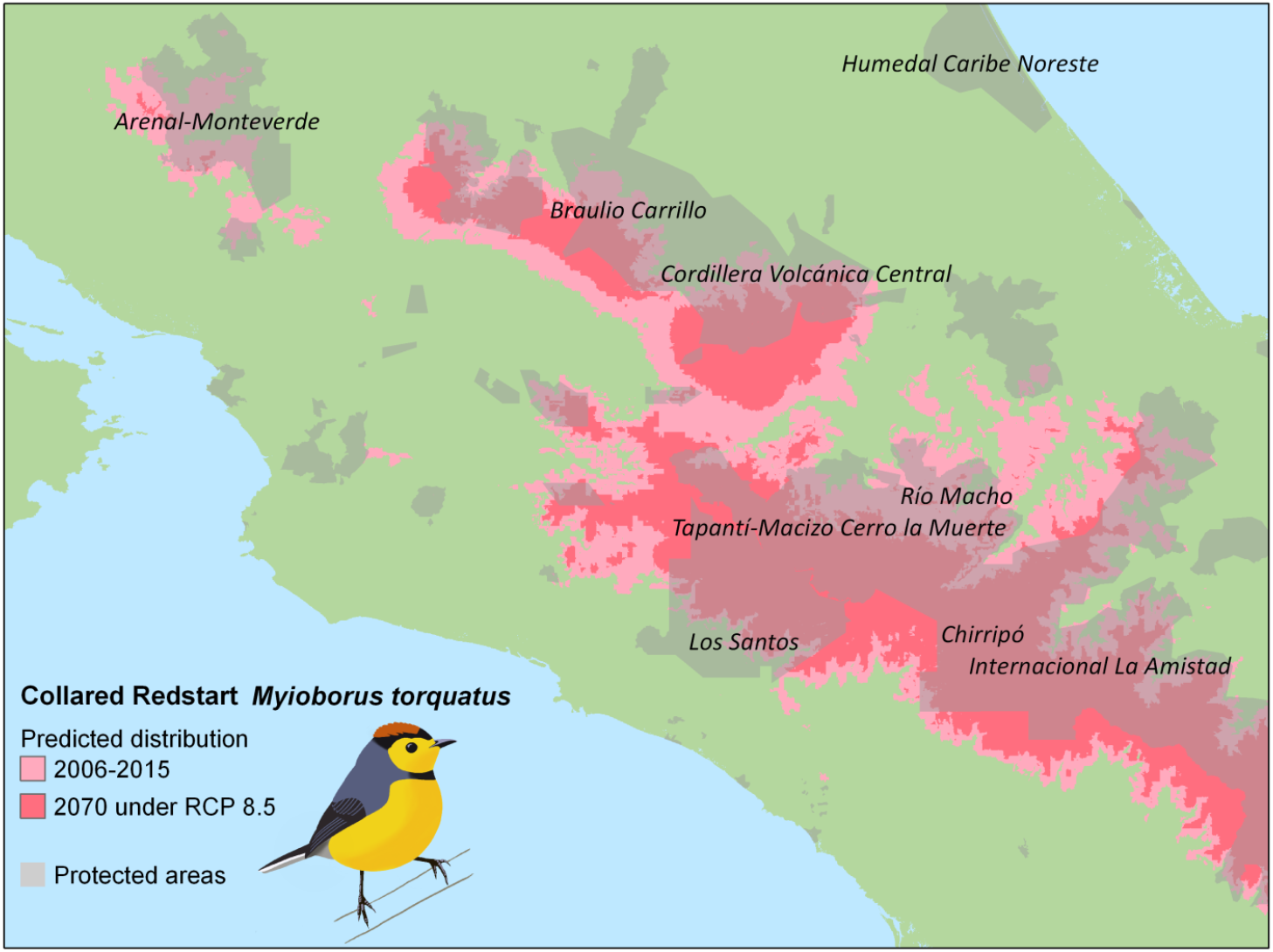
Predicted changes of Collared Redstart *Myioborus torquatus* distirbution in central Costa Rica between 206-2015 and 2070 under severe climate change (RCP 8.5), and how these changes would affect the species’ distribution within protected areas (major protected areas labelled).

**Figure 6.**
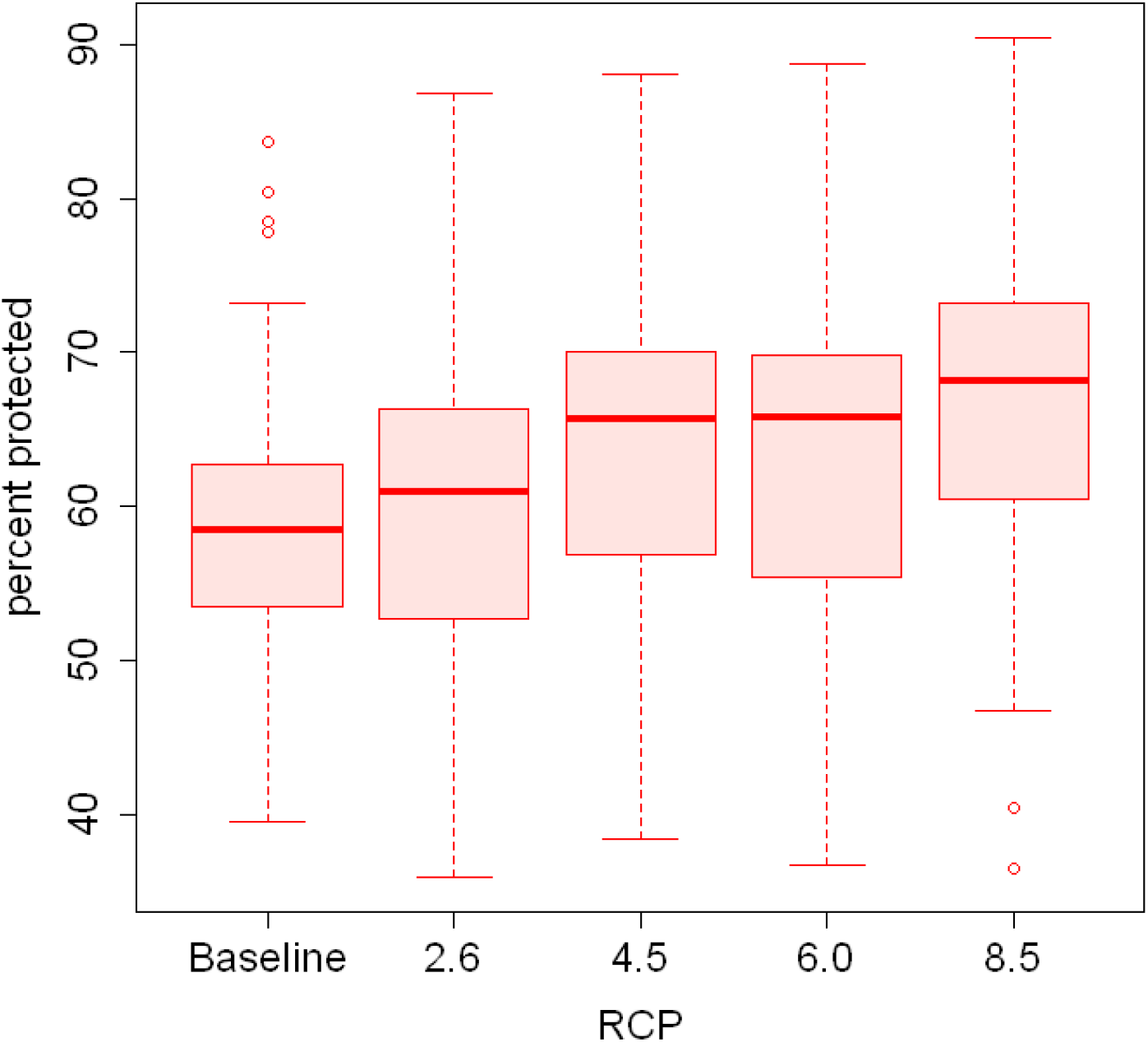
Anticipated percentage of suitable habitat within protected areas in 2006-2015 and under the four IPCC AR5 Greenhouse Gas Concentration Pathways (IPCC 2014) for 48 bird species. Circles indicate outliers.

## DISCUSSION

Our study showed that birds endemic to the Cordillera de Talamanca are likely to undergo significant range declines over the next 50 years due to climate change. The extent of those range declines could be modest, if the international community rapidly and aggressively addresses the causes of climate change (RCP 2.6) as has been proposed in the Kyoto Protocol and Paris Agreement (Oberthür and Ott 1999, Rogelj et al. 2016). But delay and inaction could push many of these species into more imperiled circumstances (other scenarios). In addition to the as yet unknown trajectory of climate action by humans, there is great uncertainty about how ecosystems will respond to climate change. If ecosystem changes are rapid, high-elevation coniferous forests and páramo shrub and high elevation grasslands could be replaced by montane forest (Kappelle and Horn 2016). The Costa Rican páramo is already small and localized, extending over just 150 km^2^, almost two-thirds of which is in one near-contiguous block around the country’s highest mountain, Cerro Chirripó (Kappelle and Horn 2016). Ecosystem replacement is therefore of particular concern for species that are endemic to páramo, such as Timberline Wren *Thryorchilus browni*, Volcano Hummingbird *Selasphorus flammula*, and Volcano Junco *Junco vulcani*, or habitat specialist as Peg-billed Finch *Acanthidops bairdii* in bamboos (*Chusquea* sp.) or the Nightingale-thrush *Catharus gracilirostris*, Sooty Thrush *Turdus nigrescens*, and Large-footed Finch *Pezopetes capitalis* in forest edges.

In addition to possible loss of habitat extent, biotic factors such as intraspecific and interspecific interactions as well as the availability of food sources, may ultimately play a significant role in species’ responses to climate change (Walther et al. 2002). For example, the Sooty Thrush is a high-elevation endemic that could be affected by the upslope spread of close relatives the Mountain Thrush *T. plebejus* and the Clay-colored Thrush *T. grayi*, which could use the same food resources and nesting sites. The extent to which high elevation bird species in the Cordillera de Talamanca are vulnerable to such factors is not known.

While our predicted average range losses were similar to those for other regions in the tropics, including Columbia (Velásquez-Tibatá et al. 2013), Bolivia and Peru (Avalos and Hernández 2015), it is important to note that unlike those studies, we did not predict that any species will lose their climactically suitable range completely. Further, unlike some other parts of the world, this region of central American has a robust network of protected areas, including the transboundary La Amistad International Park, which forms the core of the species ranges for most of the 48 species in our study (Barrantes 2009). We show that the proportion of each species’ range found within these protected areas is likely to increase, because populations are predicted to retract from the less protected lower-elevation forests. An ongoing commitment to maintain the protected areas in Costa Rica and Panama is therefore critical to the long-term prospects of the 48 bird species our study. Were habitat loss to be added to the pressures that these species face from climate change, it is quite possible that the conservation outlook for these species would be much bleaker.

Of all the species included in our study, the status Red-fronted Parrotlet is of most conservation concern. Its population size is thought to be fewer than 10,000 individuals with long-term population declines, and it is classified as Vulnerable by IUCN (BirdLife International 2004); although this assessment was based on expert opinion in the absence of data. The parrotlet’s decline is thought to be due to habitat loss on the Caribbean slop of Costa Rica where this species is found during altitudinal migrations from 3,000 down to 500 m (Stiles & Skutch 1989). Our study suggests that this species’ range losses by 2070 in the highlands will be between 30.6% and 86.5% (Table 2). Therefore, without efforts to protect the altitudinal corridor used for this species, the future of this species in the wild looks gloomy. However, this is not the only species analyzed that conducts altitudinal migrations (Table 1), and the needs of other species throughout their annual cycle need further research.

In addition to the species included in our study which are predicted to undergo rapid range contractions, there are two endemic species that we did not include in our study, due to a paucity of eBird observations. The Glow-throated Hummingbird *Selasphorus ardens* and Yellow-green Brushfinch *Atlapetes luteoviridis* are endemic to mid-elevations of the eastern Cordillera de Talamanca in Panama, and listed as Endangered and Vulnerable by the IUCN, respectively (BirdLife International 2004). Both species are thought to be vulnerable to habitat loss and fragmentation due to forest clearing in its distribution that occurred outside of protected areas (Baldwin 2020, Stiles and Sharpe 2020).

The Cordillera de Talamanca is the core range for at least 70 bird species (Barrantes 2005), even if they are not endemic to it. This list includes near endemics that have relict populations in the Darién Gap of eastern Panama and western Columbia, including the Bare-shanked Screech-Owl *Megascops clarkia* (Holt et al. 2020), and Ochraceous Wren *Troglodytes ochraceus* (Kroodsma and Brewer 2020); or endemic subspecies as Red-tailed Hawk *Buteo jamaicensis costaricensis* (Clark and Schmitt 2017)a nd Unspotted Saw-whet Owl *Aegolius ridgwayi ridgwayi* (König et al. 2009). Therefore, the habitat changes and range contractions due to climate change will affect many species beyond the 48 species included in our study, and the implications for genetic diversity of a much wider array of species merits attention.

Despite patchy occurrence data, the predicted area of occupancy in 2006-2015 closely matched the species distribution maps of BirdLife International for all of our study species, which reflects the fact that this suite of species is strongly constrained by elevation and climate (Barrantes et al. 2011). This is supported by the very high AUC values of our models, which indicates that predictions made from eBird data have high sensitivity (true negative rate) and high specificity (true negative rate). This is perhaps not surprising, because the suite of species chosen for our study are known to be climatically or elevationally restricted. However, it is known that current species range sizes are over-estimated for many species, which results in a significant under-estimation of extinction risks (Ocampo-Peñuela et al. 2016). Our study indicates that this large suite of endemic bird species will undergo range contractions, and likely, concomitant population declines in the future, but for all of them, robust population size and trend data are lacking. We recommend that a population monitoring program, possibly using a community science approach (e.g. via eBird, Kelling et al. 2019) be established to monitor trends of these species.

### Study limitations

There were some limitations in this study. The eBird data were geographically biased because there are large tracts of the Talamancan forests that are difficult to access, such as the Caribbean slope of the range. Hence, we have no way of verifying our predictions for those areas in which observations are lacking. In addition, there was a temporal mismatch between the bird data and our baseline climate data: eBird data was for 2006 to 2015, while the current climate variables were out-of-date, dating from the period 1970 to 2000. The climate data in the period of 2006-2015 were likely different from those in the period of 1970-2000 since a general warming trend was identified in the region of Central America (Aguilar et al. 2005). Moreover, our assumption which served as the foundation of the whole research was that environmental variables other than the climate are held constant until 2070 whereas in reality, this is unlikely.

While the majority of species included in our study are sedentary, six of them conduct altitudinal migrations (Table 1). We did not incorporate seasonality in our analysis, but this might be a useful future analysis, since habitat conditions in non-breeding area ranges at lower elevations could be as critical as that in their higher-elevation breeding sites. Additionally, it would be useful to widen this study to include all the non-endemic highland species that undergo elevational migration, to expand on our understanding of the effects of climate change on bird species in this region.

## CONCLUSIONS

Our analysis predicts that due to climate change, the geographic ranges of elevation-restricted birds in Costa Rica and Panama will decline substantially over the next 50 years. If this were to happen, almost all species would be candidates for upgrading from “Least Concern” to “Vulnerable”, “Endangered” or “Critically Endangered” on the IUCN Red List. However, the predicted future ranges for the majority of endemic species will be inside protected areas, which should help to lessen the likelihood of species extinctions.

## Acknowledgements

We thank Kenneth Boyle, Cody Kiefer, and Jennifer Gilmore for help with data analysis. Funding was provided by the Gettysburg College Provost’s Office.

## Appendix 1. Classification groupings of the Central American Ecosystems Map codes into 18 broad groups in our analysis

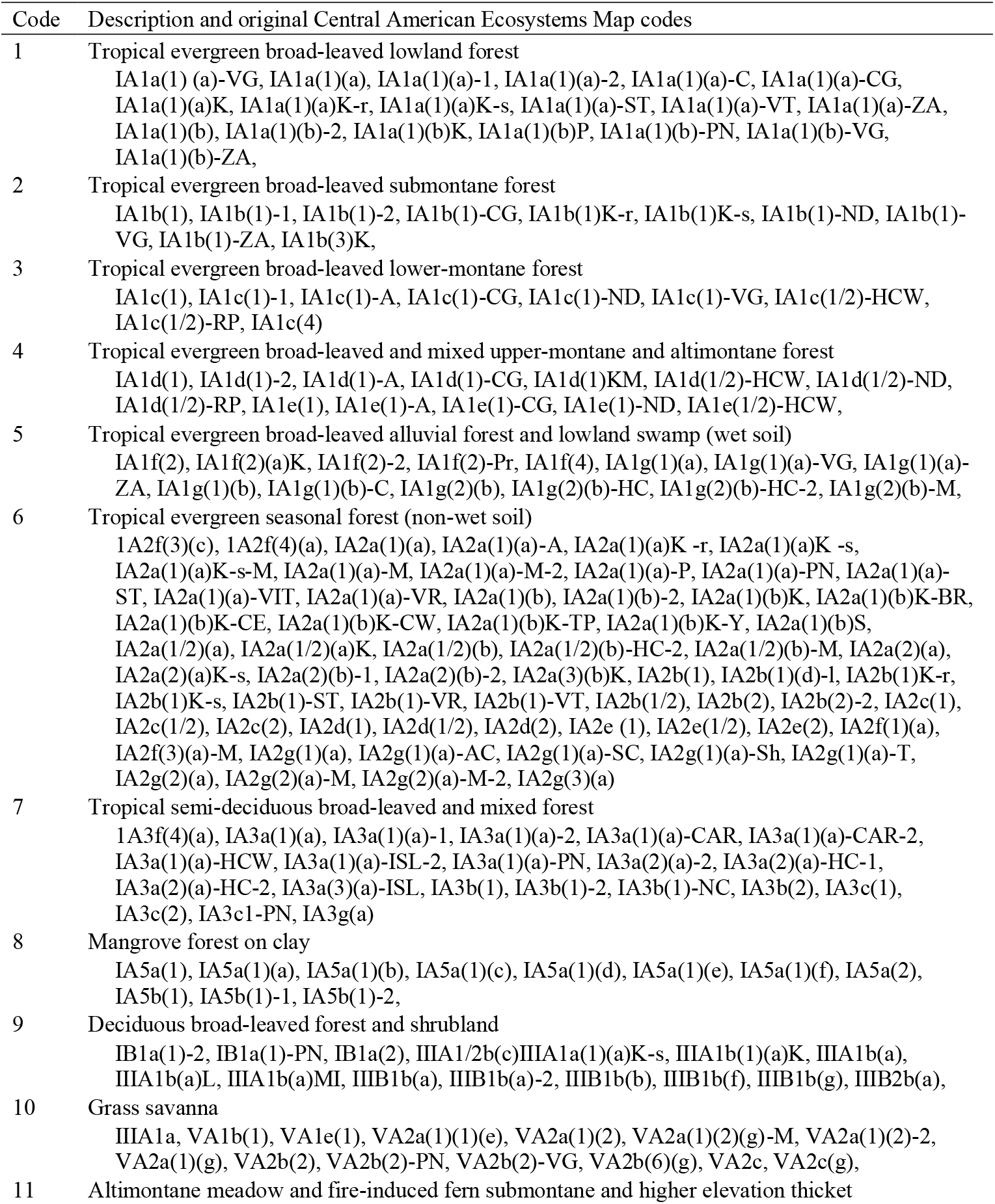
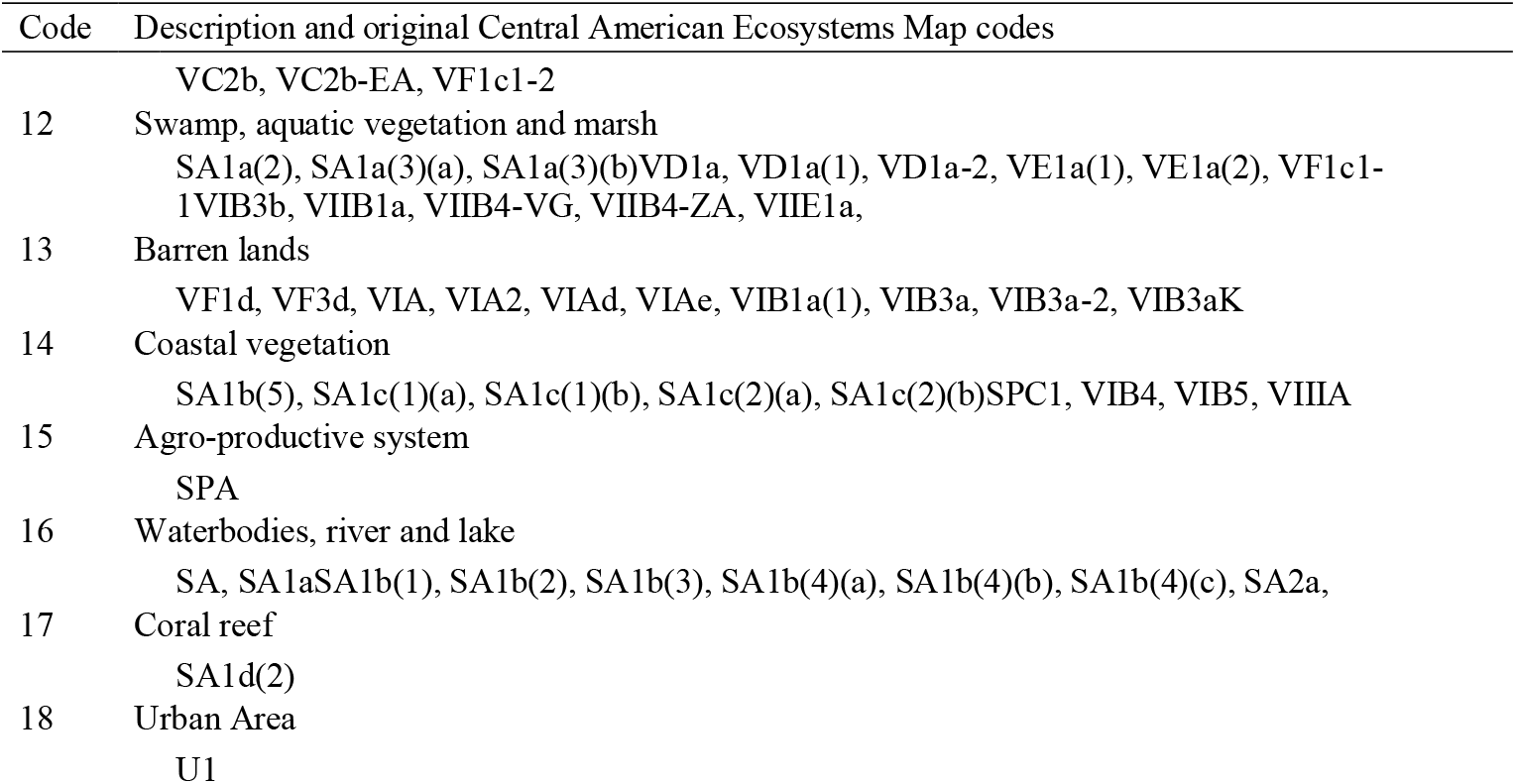

1 Appendix 1. Classification groupings of the Central American Ecosystems Map codes into 18 broad 2 groups in our analysis
| Code | Description and original Central American Ecosystems Map codes |
| --- | --- |
| 1 | Tropical evergreen broad-leaved lowland forest<br>IA1a(1) (a)-VG, IA1a(1)(a), IA1a(1)(a)-1, IA1a(1)(a)-2, IA1a(1)(a)-C, IA1a(1)(a)-CG, IA1a(1)(a)K, IA1a(1)(a)K-r, IA1a(1)(a)K-s, IA1a(1)(a)-ST, IA1a(1)(a)-VT, IA1a(1)(a)-ZA, IA1a(1)(b), IA1a(1)(b)-2, IA1a(1)(b)K, IA1a(1)(b)P, IA1a(1)(b)-PN, IA1a(1)(b)-VG, IA1a(1)(b)-ZA, |
| 2 | Tropical evergreen broad-leaved submontane forest<br>IA1b(1), IA1b(1)-1, IA1b(1)-2, IA1b(1)-CG, IA1b(1)K-r, IA1b(1)K-s, IA1b(1)-ND, IA1b(1)-VG, IA1b(1)-ZA, IA1b(3)K, |
| 3 | Tropical evergreen broad-leaved lower-montane forest<br>IA1c(1), IA1c(1)-1, IA1c(1)-A, IA1c(1)-CG, IA1c(1)-ND, IA1c(1)-VG, IA1c(1/2)-HCW, IA1c(1/2)-RP, IA1c(4) |
| 4 | Tropical evergreen broad-leaved and mixed upper-montane and altimontane forest<br>IA1d(1), IA1d(1)-2, IA1d(1)-A, IA1d(1)-CG, IA1d(1)KM, IA1d(1/2)-HCW, IA1d(1/2)-ND, IA1d(1/2)-RP, IA1e(1), IA1e(1)-A, IA1e(1)-CG, IA1e(1)-ND, IA1e(1/2)-HCW, |
| 5 | Tropical evergreen broad-leaved alluvial forest and lowland swamp (wet soil)<br>IA1f(2), IA1f(2)(a)K, IA1f(2)-2, IA1f(2)-Pr, IA1f(4), IA1g(1)(a), IA1g(1)(a)-VG, IA1g(1)(a)-ZA, IA1g(1)(b), IA1g(1)(b)-C, IA1g(2)(b), IA1g(2)(b)-HC, IA1g(2)(b)-HC-2, IA1g(2)(b)-M, |
| 6 | Tropical evergreen seasonal forest (non-wet soil)<br>IA2f(3)(c), IA2f(4)(a), IA2a(1)(a), IA2a(1)(a)-A, IA2a(1)(a)K -r, IA2a(1)(a)K -s, IA2a(1)(a)K-s-M, IA2a(1)(a)-M, IA2a(1)(a)-M-2, IA2a(1)(a)-P, IA2a(1)(a)-PN, IA2a(1)(a)-ST, IA2a(1)(a)-VIT, IA2a(1)(a)-VR, IA2a(1)(b), IA2a(1)(b)-2, IA2a(1)(b)K, IA2a(1)(b)K-BR, IA2a(1)(b)K-CE, IA2a(1)(b)K-CW, IA2a(1)(b)K-TP, IA2a(1)(b)K-Y, IA2a(1)(b)S, IA2a(1/2)(a), IA2a(1/2)(a)K, IA2a(1/2)(b), IA2a(1/2)(b)-HC-2, IA2a(1/2)(b)-M, IA2a(2)(a), IA2a(2)(a)K-s, IA2a(2)(b)-1, IA2a(2)(b)-2, IA2a(3)(b)K, IA2b(1), IA2b(1)(d)-1, IA2b(1)K-r, IA2b(1)K-s, IA2b(1)-ST, IA2b(1)-VR, IA2b(1)-VT, IA2b(1/2), IA2b(2), IA2b(2)-2, IA2c(1), IA2c(1/2), IA2c(2), IA2d(1), IA2d(1/2), IA2d(2), IA2e (1), IA2e(1/2), IA2e(2), IA2f(1)(a), IA2f(3)(a)-M, IA2g(1)(a), IA2g(1)(a)-AC, IA2g(1)(a)-SC, IA2g(1)(a)-Sh, IA2g(1)(a)-T, IA2g(2)(a), IA2g(2)(a)-M, IA2g(2)(a)-M-2, IA2g(3)(a) |
| 7 | Tropical semi-deciduous broad-leaved and mixed forest<br>IA3f(4)(a), IA3a(1)(a), IA3a(1)(a)-1, IA3a(1)(a)-2, IA3a(1)(a)-CAR, IA3a(1)(a)-CAR-2, IA3a(1)(a)-HCW, IA3a(1)(a)-ISL-2, IA3a(1)(a)-PN, IA3a(2)(a)-2, IA3a(2)(a)-HC-1, IA3a(2)(a)-HC-2, IA3a(3)(a)-ISL, IA3b(1), IA3b(1)-2, IA3b(1)-NC, IA3b(2), IA3c(1), IA3c(2), IA3c1-PN, IA3g(a) |
| 8 | Mangrove forest on clay<br>IA5a(1), IA5a(1)(a), IA5a(1)(b), IA5a(1)(c), IA5a(1)(d), IA5a(1)(e), IA5a(1)(f), IA5a(2), IA5b(1), IA5b(1)-1, IA5b(1)-2, |
| 9 | Deciduous broad-leaved forest and shrubland<br>IB1a(1)-2, IB1a(1)-PN, IB1a(2), IIIA1/2b(c)IIIA1a(1)(a)K-s, IIIA1b(1)(a)K, IIIA1b(a), IIIA1b(a)L, IIIA1b(a)MI, IIIB1b(a), IIIB1b(a)-2, IIIB1b(b), IIIB1b(f), IIIB1b(g), IIIB2b(a), |
| 10 | Grass savanna<br>IIIA1a, VA1b(1), VA1e(1), VA2a(1)(1)(e), VA2a(1)(2), VA2a(1)(2)(g)-M, VA2a(1)(2)-2, VA2a(1)(g), VA2b(2), VA2b(2)-PN, VA2b(2)-VG, VA2b(6)(g), VA2c, VA2c(g), |
| 11 | Altimontane meadow and fire-induced fern submontane and higher elevation thicket |
|  | VC2b, VC2b-EA, VF1c1-2 |
| 12 | Swamp, aquatic vegetation and marsh<br>SA1a(2), SA1a(3)(a), SA1a(3)(b)VD1a, VD1a(1), VD1a-2, VE1a(1), VE1a(2), VF1c1-1VIB3b, VIIB1a, VIIB4-VG, VIIB4-ZA, VIIE1a, |
| 13 | Barren lands<br>VF1d, VF3d, VIA, VIA2, VIAd, VIAe, VIB1a(1), VIB3a, VIB3a-2, VIB3aK |
| 14 | Coastal vegetation<br>SA1b(5), SA1c(1)(a), SA1c(1)(b), SA1c(2)(a), SA1c(2)(b)SPC1, VIB4, VIB5, VIIIA |
| 15 | Agro-productive system<br>SPA |
| 16 | Waterbodies, river and lake<br>SA, SA1aSA1b(1), SA1b(2), SA1b(3), SA1b(4)(a), SA1b(4)(b), SA1b(4)(c), SA2a, |
| 17 | Coral reef<br>SA1d(2) |
| 18 | Urban Area<br>U1 |

